# Indomethacin-induced gut damage in a surrogate insect model, *Galleria mellonella*

**DOI:** 10.1101/606319

**Authors:** Helena Emery, Richard Johnston, Andrew F. Rowley, Christopher J. Coates

**Author notes:** Corresponding author: C.J. Coates, PhD.

## Abstract

Indomethacin is a non-steroidal anti-inflammatory drug that causes gastric ulceration and increased ‘leakiness’ in rat models, and is used routinely as an assay to screen novel compounds for repair and restitution properties. We set out to establish conditions for indomethacin-induced gut damage in wax-moth (*Galleria mellonella*) larvae with a view to reducing the need for rodents in such experimentation. We administered indomethacin (1 – 7.5 μg/larva) to *G. mellonella* via intrahaemocoelic injection and gavage (force-feeding) and monitored larval survival and development, blood cell (haemocyte) numbers, and changes in gut permeability. Increased levels of gut leakiness were observed within the first 4 to 24-hours by tracking fluorescent microspheres in the faeces and haemolymph (blood equivalent). Additionally, we recorded varying levels of tissue damage in histological sections of the insect midgut, including epithelial sloughing and cell necrosis. Degeneration of the midgut was accompanied by significant increases in detoxification-associated activities (superoxide dismutase and glutathione-S-transferase). Herein, we present the first evidence that *G. mellonella* larvae force-fed indomethacin display broad symptoms of gastric damage similar to rodent models.

## Introduction

When considering more carefully our use of vertebrates in experimentation, there is constant need to develop alternative model systems *in vitro*, *in vivo* or *in silico*. One such *in vivo* alternative is the larvae of the greater wax-moth, *Galleria mellonella*. These insects are now used widespread as ‘mini-hosts’ for the investigation of microbial pathogenicity (Mowlds *et al.*, 2010; Kloezen *et al.*, 2015; Lim *et al.*, 2018; Cools *et al.*, 2019), screening of xenobiotics/toxins (Allegra *et al.*, 2018; Coates *et al.*, 2018) and functional characterisation of virulence factors (Altincicek *et al.*, 2007; Champion *et al.*, 2016). Larvae of *G. mellonella* are designated a non-animal technology (Allegra *et al.*, 2018), which means there are fewer ethical restrictions and regulations compared to vertebrates. Additionally, practical advantages include low maintenance costs, thermal tolerance to 37°C, ease of use (accurate dosages) and high turnover – results can be obtained within 72 hours – in contrast to vertebrates. Although *G. mellonella* have been used to study the infectivity of gut pathogens such as *Campylobacter jejuni* (Senior *et al.*, 2011), *Listeria monocytogenes* (Mukherjee *et al.*, 2013a), *Vibrio* spp. Wagley *et al.*, 2018), *Shigella* spp. (Barnoy *et al.*, 2017) and *Salmonella enterica* (Card *et al.*, 2016), there remains a distinct lack of information regarding the alimentary canal of this insect, and its role in pathogenesis.

From mouth to rectum, the digestive system of lepidopteran larvae (like *G. mellonella*) consists of three distinct regions: the foregut (stomatodaeum), midgut (mesenteron) and hindgut (proctodaeum) (Engel and Moran, 2013; Linser and Dinglasan, 2014). The midgut is the primary site of digestion and absorption in many insects; it lacks the exoskeletal/chitin lining seen in the fore- and hind-guts. The basic tissue architecture of the midgut is similar to those found in the human intestine, such as epithelial arrangements of columnar cells and smooth septate junctions that control permeability – analogous to tight junctions (Green *et al.*, 1980). The insect peritrophic matrix is the functional equivalent to the mammalian mucus layer, which acts as a barrier for the epithelial cells and impedes pathogen movement into the body cavity (i.e., the haemocoel) (Campbell *et al.*, 2008; Kuraishi *et al.*, 2011). Moreover, some microbial communities characterised in the midgut of *G. mellonella* are similar to those found in crypts of the human intestine (Mukherjee *et al.*, 2013b).

Non-steroidal anti-inflammatory drugs (NSAIDs) are used widespread to alleviate pain by inhibiting the activities of cycloxygenase isozymes (COX1, COX2), yet known side effects manifest in gastrointestinal injury (Playford *et al.*, 1999; Brune and Patrignani, 2015). In particular, indomethacin causes ulceration by inducing apoptosis, reducing gastric blood flow and activating innate immune cells (neutrophils), which all contribute to mucosal secretion, maintenance and defence (Marchbank *et al.*, 2011; Matsui *et al.*, 2011). Gastric complications arising from indoemthacin exposure have been developed into a standard rodent restraint/ulcer assay in order to screen novel compounds and health food supplements for putative therapeutic properties – tissue repair and restitution (Playford *et al.*, 1999 and 2001; Mahmood *et al.*, 2007). The adverse effects of indomethacin, notably inflammation, permeability and REDOX imbalance, have been studied thoroughly in humans and rodent models (Basivireddy *et al.*, 2003; Bjarnason and Takeuchi, 2009; Sigthorsson *et al.*, 2000; Perron *et al.*, 2013). Therefore, we set out to interrogate the putative effects of indomethacin on the alimentary canal of *G. mellonella*. Firstly, we utilised two inoculation methods (intrahaemocoelic injection and gavage) to assess the relative toxicity of indomethacin in insect larvae across the concentration range, 0 – 7.5 μg/larva. Secondly, we combined histopathology screening, X-ray microtomography/microscopy (XRM) and enzyme activity to assess the integrity of the midgut tissues in the absence and presence of this NSAID. Indomethacin treatments were established by extrapolating from Marchbank *et al.*, (2011), wherein a dose of 20 mg/kg was administered to induce gastric damage in rats – the equivalent being 5.0 μg/larva.

## Materials and Methods

### Reagents

Unless stated otherwise, all chemicals and reagents were sourced from Sigma Aldrich (Dorset UK) in their purest form listed. Green and blue fluorescent (latex) microspheres ranging from 0.5 to 6 μm in diameter were purchased from Polysciences, Inc. (Fluoresbite®). Stock solutions of indomethacin were prepared in 5% dimethyl sulfoxide (DMSO; v/v) and diluted in filter-sterilized (0.2 μm) phosphate buffered saline (PBS) pH 7.4.

### Insects

Larvae of *G. mellonella* (final instar) were purchased from Livefoods Direct Ltd (Sheffield UK). Upon arrival, each larva was inspected for signs of damage, infection and melanisation. Healthy larvae weighing between 0.25 g and 0.35g were retained and stored at 15°C in the dark for no more than 7 days.

### Larval survival and pupation studies

Larvae were assigned randomly to each treatment (*n* = 10 per replicate) and placed in 90 mm petri dishes lined with Whatman filter paper and wood shavings from the commercial supplier. Indomethacin was administered by intrahaemocoelic injection (INJ) or force feeding (FF; gavage) using a sterile 27-gauge hypodermic needle across the concentration range, 0 to 7.5 μg/larva. The volume of each inoculum was standardised to 10 μL. Negative controls consisted of PBS pH 7.4 alone and PBS containing 5% dimethyl sulfoxide (DMSO). Post-inoculation, larvae were incubated at 30°C and assessed for mortality (response to prodding) and development (pupation events) for up to 10 days.

### Haemocyte counts and viability

Larvae treated with indomethacin, PBS, or 5% DMSO were assessed for haemocyte numbers within 4 and 72 hours post-inoculation. Insects were chilled on ice for ~3 minutes prior to injection of 100 μL anticoagulant (15 mM NaCl, 155 mM trisodium citrate, 0.03 mM citric acid, 0.02 mM disodium EDTA). Larvae containing anti-coagulant were placed back on ice for 2 minutes prior to piercing the integument above the head using a 27-gauge hypodermic needle. Haemolymph was collected into pre-chilled, sterile microcentrifuge tubes. For haemocyte viability, haemolymph was extracted in the absence of anticoagulant and mixed in a ratio of 1:5 with 0.4% trypan blue (w/v in PBS) and incubated at room temperature for 1 to 2 minutes. In all cases, haemolymph samples were applied to an improved Neubauer haemocytometer for cell enumeration using bright-field optics of a compound microscope. Two technical replicates were performed per sample. Haemocytes were considered dead when the cytoplasm stained positively for trypan-blue (Strober, 2015).

### Detoxification-associated assays

Haemolymph was extracted (as stated above) and pooled from three larvae at 4, 24, 48 and 72 hours after being force-fed indomethacin. Haemolymph was centrifuged at 500 × g for 5 minutes at 4°C to pellet the haemocytes. Approximately, 2.5 ×10^5^ haemocytes from each treatment/time point were added to lysis buffer (50mM potassium phosphate, 2mM EDTA, 1mM DTT and a proteinase inhibitor cocktail (Roche cOMPLETE™ Mini kit) and centrifuged at 14,000 × *g* for 10 minutes at 4°C. Supernatants were retained and stored at −80°C. Midgut tissues from three larvae per treatment/time point were dissected out and washed in PBS. Approximately, 30 mg of tissue was placed in 1 ml lysis buffer and homogenised prior to centrifugation and storage as described above. Protein concentrations of haemolymph and midgut samples were determined using the Biuret method with bovine serum albumin (BSA) as a standard.

#### a. Superoxide dismutase (SOD) activity

SOD activity (EC 1.15.1.1) was determined by using the method described by Dubovskiy *et al.* (2008). Briefly, 80 μL of sample (haemolymph or homogenised midgut) was mixed with reaction solution (500 μL PBS containing 70 μM NBT, 125 μM xanthine), followed by the addition of 20 μL xanthine oxidase solution (5 mg BSA, 15 μL xanthine oxidase (20 units/ml) per ml PBS), and incubated for 20 minutes at 28°C. Total assay volume was 600 μL. Xanthine oxidase catalyses xanthine to produce superoxide anions (O_2_^−^), which reduce NBT to a formazan dye. The inhibition of NBT reduction is indicative of SOD activity, which is monitored spectrophotometrically at 560 nm. SOD activity is presented as the increase in absorbance (560 nm) per min per mg protein.

#### b. Glutathione S-transferase (GST) activity

GST activity (EC 2.5.1.18) was determined using the method described by Dubovskiy *et al.* (2008). Briely, 20 μL of sample was mixed with 500 μL GST assay solution (1 mM glutathione, 1mM 1-chloro-2,4-dinitrobenzene (DNCB) dissolved in PBS) and incubated at 28 °C for 5 minutes. The reaction of DNCB and glutathione is catalysed by GST, producing 5-(2,4-dinitrophenyl)-glutathione – detectable at 340 nm. GST activity is presented as the increase in absorbance (340 nm) per min per mg protein.

### Gut permeability assessments

Fluorescently tagged, carboxylate-modified latex microspheres (1 ×10^6^) of 0.5 μm, 1 μm, 2 μm and 6 μm in diameter were resuspended in 10 μL PBS (control) or 10 μL indomethacin solution (1 or 7.5 μg dose) in order to form co-inoculates (thereby avoiding piercing an insect twice). Larvae were force-fed 10 μL of each co-inoculate and incubated at 30°C until haemolymph and faeces were collected from each treatment group at 4, 24, 48 and 72 hours. Faeces were homogenised in 1 ml PBS pH 7.4. The number of microspheres in the haemolymph/faeces were enumerated using a fluorescent microscope (Olympus B×43f).

### Histopathology of the insect alimentary canal

Larvae force-fed indomethacin (7.5 μg per insect) or PBS (negative control) were sacrificed at 4, 24, 48 and 72 hours post-inoculation by intrahaemocoelic injection of 100 μL 10% buffered formalin, immediately prior to submersion in the same solution for 24 hours. Larvae were cut into 3 sections, head, middle and posterior (anus), and stored in 70% ethanol prior to wax embedding. Briefly, each sample was dehydrated using an ethanol series, 70%, 80% and 90% for one hour each, followed by 3x 1 hour 100% ethanol washes. Dehydrated samples were washed twice in HistoClear or HistoChoice (Sigma Aldrich) for 1 hour each to remove any remaining fixative. Samples were resuspended in 50:50 HistoChoice : parafin wax for 1 hour prior to complete wax embedding. Embedded samples were cut into ~ 6 μm sections using a microtome, adhered to glass slides with egg albumin solution (~1% w/v), and dried for 24 hours before staining. Loaded slides were stained using Cole’s hematoxylin and eosin (see Supp. Materials/Materials for further details).

### X-ray microtomography/microscopy of *Galleria mellonella*

Larvae force-fed PBS were sacrificed at 72 hours and fixed with 10% formalin for 30 hours, followed by dehydration in 100% ethanol for 24 hours. Samples were submerged in Lugol’s iodine (PRO.LAB Diagnostics) for 2 weeks and washed with 70% ethanol prior to microscopy. Insect samples were analysed via X-ray microscopy (XRM) using a lab-based Zeiss Xradia 520 (Carl Zeiss XRM, Pleasanton, CA, USA) X-ray Microscope attached to a CCD detector system with scintillator-coupled visible light optics, and tungsten transmission target. The specimen was placed in a plastic screw-top microcentrifuge tube and submerged in 75% ethanol to prevent the tissues from drying out. To achieve a higher resolution over the entire organism, the insect was imaged along its ~25 mm length at high resolution, using an overlap-scan and stitching procedure including five individual scans, with 44% overlap between each scan. An X-ray tube voltage of 80 kV, and a tube current of 87 μA were used, with an exposure of 1000 ms, and a total of 3201 projections. An objective lens giving an optical magnification of 0.4 was selected with binning set to 2, producing an isotropic voxel (3-D pixel) size of 8.5635 μm. The individual tomograms were reconstructed from 2-D projections and stitched into a single large volume using a Zeiss commercial software package (XMReconstructor, Carl Zeiss), utilising a cone-beam reconstruction algorithm based on filtered back-projection. XMReconstructor was also used to produce 2-D grey scale slices for subsequent analysis. Drishti and Drishti Paint software V2.6.4. (Limaye, 2012) was employed to highlight regions of interest and digitally segment the alimentary canal.

### Data handling

All data were gathered from experiments carried out on at least three independent occasions and are represented by mean values ± standard error (see individual figure legends for sample sizes). D’Agostino and Pearson normality tests were performed, and if necessary, data were square-root transformed. Results from survival studies were analysed using the log-rank (Mantel-Cox) test for comparing curves, whereas larval pupation levels, total haemocyte counts, cell deaths, and faecal/haemolymph micropshere loads were analysed using 2- or 3-way ANOVA with Tukey’s multiple comparison tests in GraphPad Prism v7. A Pearson’s Correlation test was employed to assess the relationship between microsphere size (0.5 – 6 μm) and their presence in the haemolymph of control larvae. Enzyme assays, GST and SOD, were analysed in ‘R’ studio using General Liner Hypotheses (ghlt). In all cases, differences were considered significant when P ≤ 0.05.

Histology slides were single-blind assessed using paired treatment (n = 13) and control (n = 13) samples from 4 to 72 hours. A grading system (1 – 4) was used to catagorise the extent of tissue damage(s) or lack thereof (Supp. Materials/Materials for descriptions). Drishti and Drishti Paint V2.6.4. (Limaye, 2012) and ImageJ (Abrámoff *et al.*, 2004) softwares were used to process/present microscopy findings. Images were adjusted for colour balance and contrast/brightness only.

## Results

### Evaluating the relative toxicity of indomethacin on Galleria mellonella

#### a) Survival

Intrahaemocoelic injection of indomethacin had no measurable negative impacts on larval survival across the concentration range 0.5–7.5 μg/larva (0% mortality), which is the equivalent to 2–30 mg/kg in rodents. Oral administration of indomethacin (force-feeding) in excess of 2.5 μg/larva led to a 7–10% decline in survival within 72 hours (Figure 1), but was not found to be statistically significant overall (log-rank (Mantel-Cox) test; *X*^*2*^(6) = 11.71, *P* = 0.0688). Further inspection of the survival curves revealed no statistical differences between the PBS/DMSO controls and either 2.5 μg/larva (*X*^*2*^(1) = 2.034, P = 0.1538) or 5–7.5 μg/larva (*X*^*2*^(1) = 3.105, P = 0.0780).

**Figure 1.**
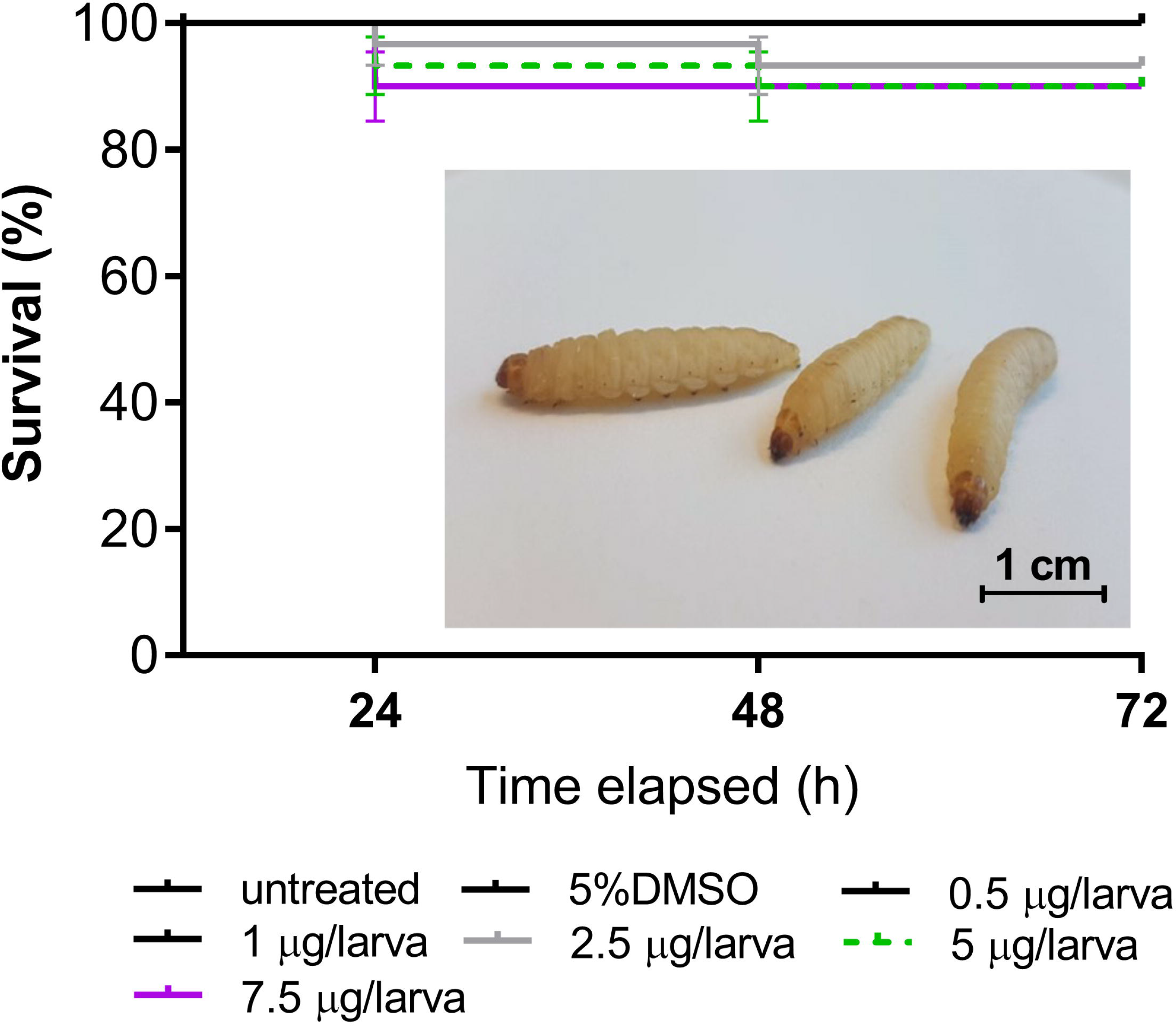
Survival of *Galleria mellonella* larvae following force-feeding (gavage) of indomethacin, 0.5 – 7.5 μg/larva. Post-inoculation, larvae were maintaned in darkness at 30°C for 72 hours. Larvae that were unresponsive to being rolled over or prodded were considered dead. Values are expressed as means ± SE (*n* = 30 per treatment, 210 in total). Inset: three healthy (untreated) *G. mellonella* larvae.

#### b) Development

Approximately 87% of untreated larvae transitioned into pupae after 6 days incubation at 30°C (experimental temperature) – increasing to 97% by day 10 (Figure 2). No more than 40% of *G. mellonella* force-fed indomethacin (0.5–7.5 μg/larva) or 5% DMSO (absent indomethacin) were observed pupating at day 6. By day 10, numbers of pupae increased to 80-90% for injected larvae (Figure 2a) and ≤63% for force-fed larvae (Figure 2b). Using a three-way ANOVA, we determined time (Days 6 and 10) and inoculation method (FF and INJ) to account for 38% (*F*(1, 24) = 50.02, P < 0.0001) and 41% (*F*(1, 24) = 54.19, P < 0.0001) of the variation, respectively. Less than 0.5% of the variation within the data can be attributed to treatment: 5% DMSO (negative control), 1 μg (low dose) and 7.5 μg (high dose) indomethacin per larva (*F*(2, 24) = 0.0833, P = 0.92). Tukey’s multiple comparison (post-hoc) tests revealed statistical differences between the untreated insects and those force-fed indomethacin or 5% DMSO only (Figure 2b), with no differences detected between mean values of injected insects at any time-point. These data suggest the route of administration alone can delay the onset of pupation, and it is unlikely due to indomethacin exposure.

**Figure 2.**
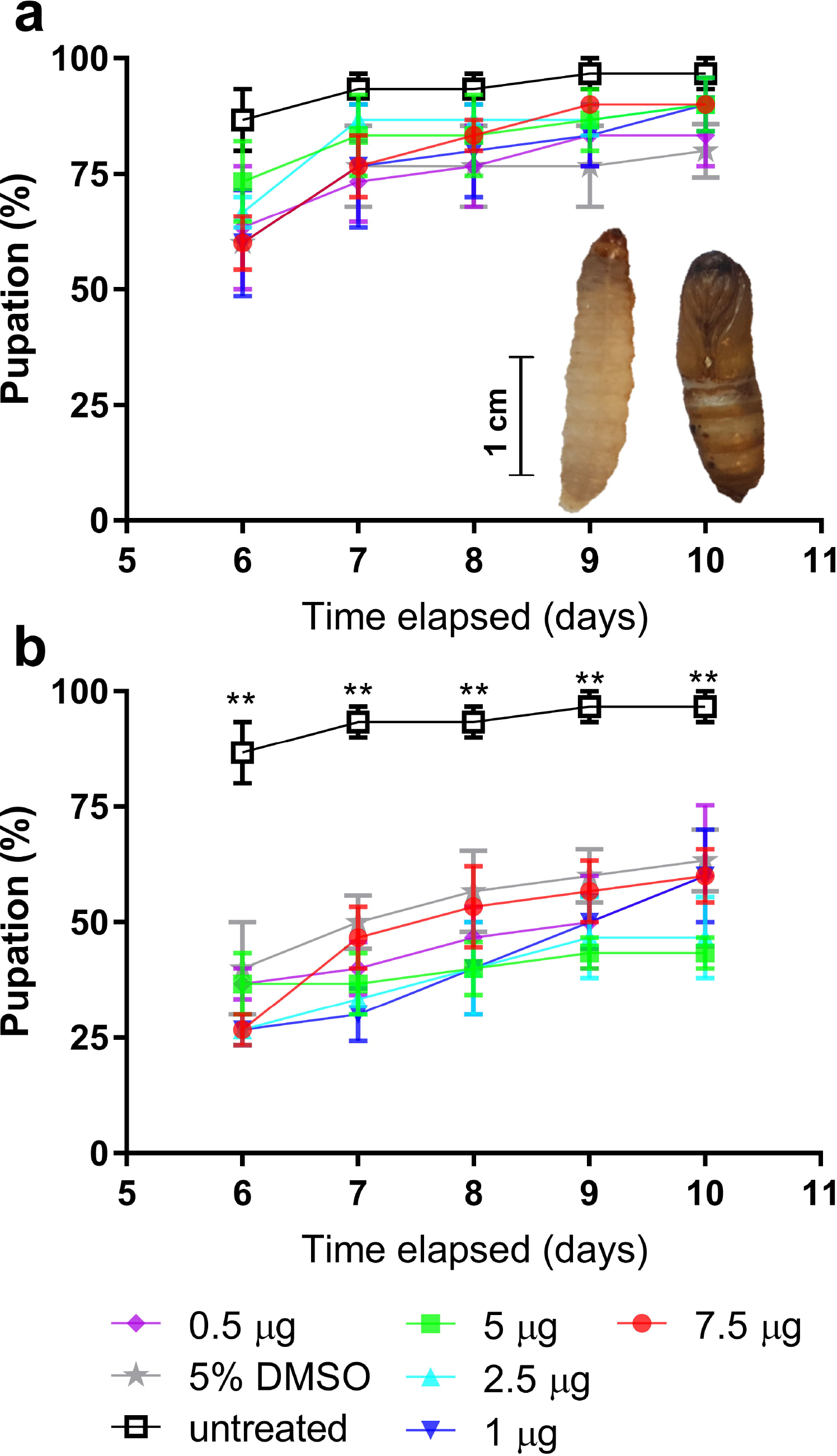
Development (pupation) of *Galleria mellonella* larvae following inoculation with indomethacin, 0.5 – 7.5 μg/larva. Larvae received indomethacin via intrahaemocoelic injection (**a**) or force feeding (**b**), and were maintaned subsequently in darkness at 30°C for 10 days. The number of larve undergoing pupation was recorded. Inset (a): typical appearances of a healthy larva (left image) and a moth pupa (right image). Values are expressed as means ± SE (*n* = 30 per treatment, 390 in total across both inoculation methods). Symbol: ** = P < 0.01 when comparing untreated (negative controls) to all other treatments at the respective time points.

#### c) Circulating blood cells (haemocytes)

To investigate further any potential non-target effects of indomethacin, we monitored immune cell (haemocyte) numbers and levels of cell death within the insect haemolymph (i.e., blood). Total haemocyte counts varied little in untreated and PBS control insects over the 72-hour experimental period, 2.5 – 3 ×10^6^ mL^−1^ and 2.7 – 3.7 ×10^6^; mL^−1^, respectively (Figure 3). At 24 hours post-inoculation, all concentrations of indomethacin injected into the haemolymph led to substantial increases in haemocyte numbers (up to 5 ×10^6^ mL^−1^) compared to the untreated/controls(<2.8 ×10^6^ mL^−1^; Figure 3a). Force-feeding indomethacin led to increased circulating haemocyte numbers at the highest dose of 7.5 μg/larva only (4.4 ×10^6^ mL^−1^; Figure 3b). In all cases, haemocyte numbers returned to control levels by 48 hours. A 2-way ANOVA revealed treatment to account for 15–19% of the variation within the data (INJ - *F*(4, 40) = 4.488, P = 0.0043; FF - *F*(4, 40) = 2.988, P = 0.0300). Time accounted for <5% of the variation in force-fed insects (Time, *F*(3, 40) = 1.033, P = 0.3884) but ~30% for injected insects (Time, *F*(3, 40) = 11.84, P < 0.0001). These data likely reflect the unimpeded exposure of haemocytes to indomethacin when it is administered directly into the haemocoel (body cavity), whereas the gut presents a natural barrier when administered via force-feeding.

**Figure 3.**
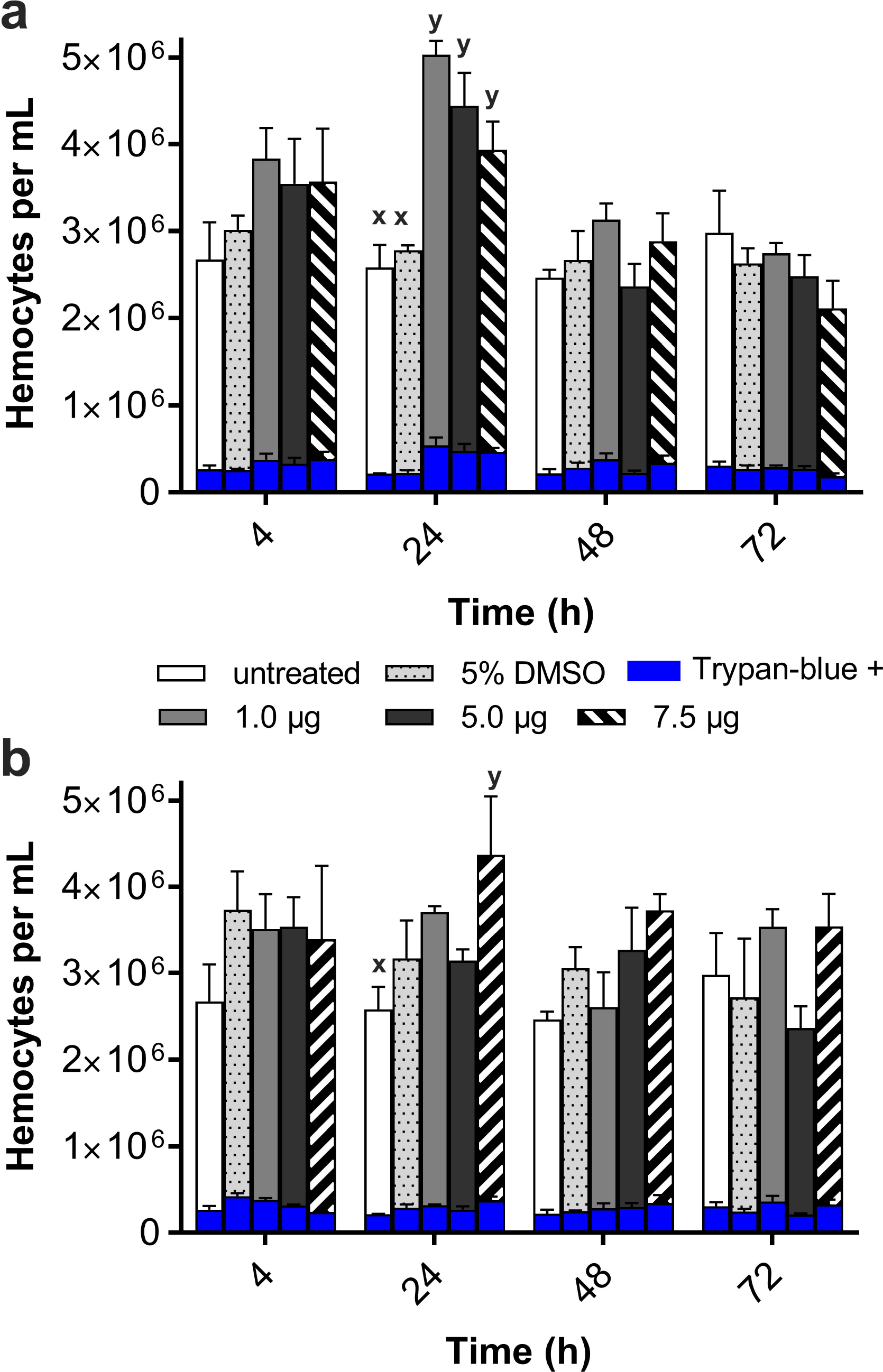
Haemocyte responses of *Galleria mellonella* larvae following inoculation of indomethacin, 1 – 7.5 μg/larva. Larvae received indomethacin via intrahaemocoelic injection (**a**) or force feeding (**b**), and were maintaned subsequently in darkness at 30°C for 3 days. Total haemocyte counts were performed, and haemocyte viability was determined using the trypan blue exclusion assay (dead cell numbers are represented by the blue bars in panels **a** and **b**). Values are presented as the mean ± SE (n = 36 per treatment, 324 in total across both inoculation methods). Unshared letters (x, y) represent significant differences (P ≤ 0.05) determined by Tukey’s multiple comparison tests.

Concerning cytotoxicity, proportions of haemocytes staining positively for trypan-blue (i.e., dead or dying) ranged from 7–12% for the duration of the experiment – regardless of the inoculation method used or dose of indomethacin (Figure 3; Supp. Table 1).

### Characterising the effects of indomethacin on the midgut of Galleria mellonella

#### a) Alimentary canal mapping

Using X-ray microtomography, we mapped out the alimentary canal and integumentary musculature of *G. mellonella* larvae (n = 3) over their entire ~25 mm lengths with a resolution of 8.6 μm (Figures 4a - 4c). Like most insects, the alimentary canal can be sub-categorised into three regions: foregut, midgut and hindgut (Figure 4b). The midgut is disinguishable from the fore– and hind-guts due to the absence of a cuticle lining, and our measurements indicate that the midgut tissues make-up >50% of the alimentary canal (n = 3). The midgut is arranged into pleats or folds of columnar epithelial cells and goblet cells that produce and maintain the peritrophic matrix (functional equivalent to the human mucus layer). Visceral (striated) muscles surround the gut tissues (anchoring the cellular arrangements in place), and represent the final layer between the gut contents and the haemolymph.

**Figure 4.**
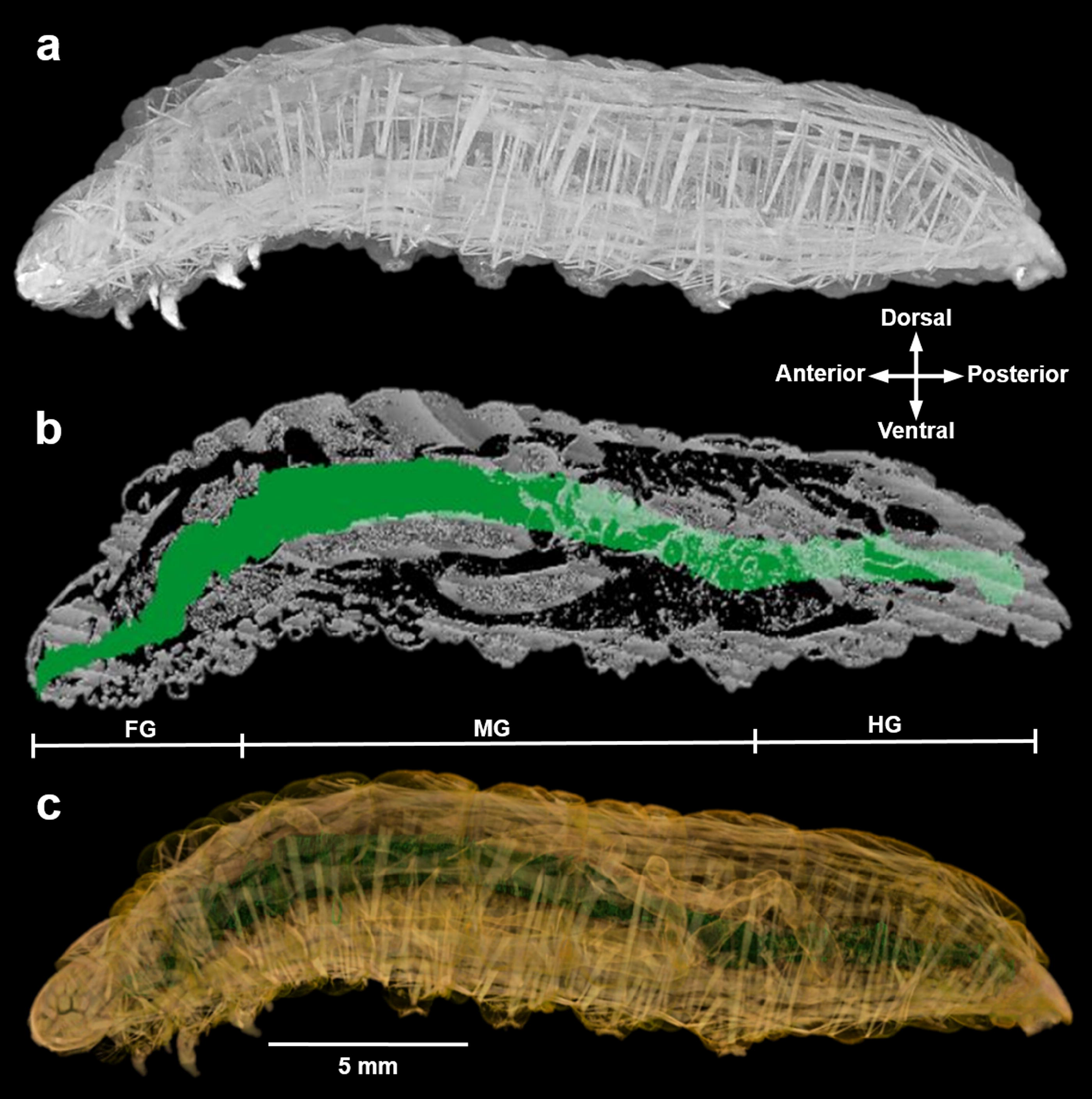
X-ray microtomography of *Galleria mellonella.* **a**) 3-dimensional render of a representative insect larva stained with Lugol’s iodine. Striated muscle fibres are distributed throughout the insect – muscles are attached to the integument and run through the cuticle layers. **b**) The entire alimentary canal of the insect larva was highlighted by manually inspecting and colouring 1014 slices using Drishti software (Limaye, 2012). FG, foregut; MG, midgut; HD, hindgut. **c**) Combined 3-dimensional render with the integument architecture coloured yellow and the alimentary canal coloured green.

#### b) Gut permeability

Indomethacin and NSAIDs broadly increase gastrointestinal ‘leakiness’ in humans and rodents, with tissue damage located in the small intestine and colon (Smecuol *et al.*, 2001). In order to test whether indomethacin caused similar pathological symptoms in the insect alimentary canal, we force-fed *G. mellonella* inocula containing indomethacin (0, 1 or 7 μg/larva) and a selection of microspheres, 0.5 – 6 μm in diameter (Figure 5). In the absence of indomethacin (PBS control), the majority of 6 μm and 2 μm spheres (36 – 58%) made their way down the alimentary canal and were defecated within 24 hours (Figure 5a and 5b), whereas 1 μm spheres (33%) were released later at 48 hours (Figure 5c). The presence of indomethacin (1 or 7.5 μg/larva) led to substantial increases in the number of microspheres (0.5 – 6 μm) detected in the haemolymph (blood) and concomitantly fewer were recovered from faeces. At 4 and 24 hours post-indomethacin treatment, 27–31 ×10^4^ (6 μm) spheres leaked from the gut into the haemolymph in comparison to the control, 10–14 ×10^4^(Figure 5a). Notably, the presence of microspheres in the haemolymph was recognised by phagocytic haemocytes and subsequently internalised (Supp. Figure 1). Independent of microsphere size, indomethacin exacerbated gut leakiness by 1.4 to 3-fold: 6 μm (*F*(2, 24) = 5.801, P = 0.0042), 2 μm (*F*(2, 24) = 11.04, P < 0.0001), 1 μm (*F*(2, 24) = 21.48, P < 0.0001), and 0.5 μm (*F*(2, 24) = 22.75, P < 0.0001), and accounted for 9–30% of the variation in haemolymph loads. In contrast, time accounted for the majority of variation, 25–51%, in faecal microsphere loads: 6 μm (*F*(3, 24) = 17.38, P < 0.0001), 2 μm (*F*(3, 24) = 3.57, P = 0.0288), 1 μm (*F*(3, 24) = 3.94, P = 0.0204), and 0.5 μm (*F*(3, 24) = 8.378, P = 0.0006).

**Figure 5.**
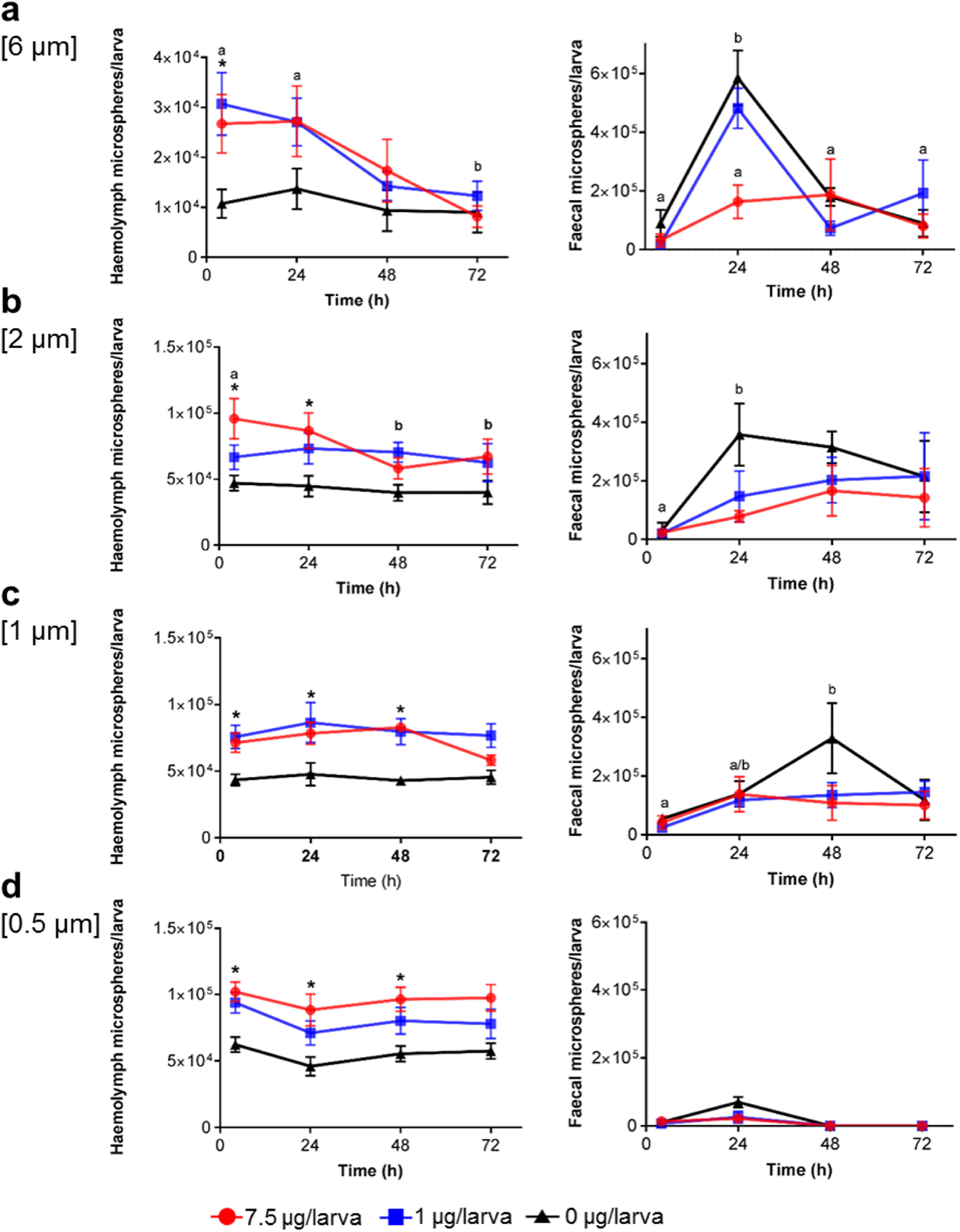
Gut permeability of *Galleria mellonella* larvae following force feeding of indomethacin, 0 – 7.5 μg/larva. Permeability (or leakiness) was determined by the number of 6 μm (a), 2 μm (b), 1 μm (c) and 0.5 μm (d) spheres found in the haemolymph and faeces between 4 and 72 hours post-inoculation. Each larva was co-inoculated with 1 ×10^6^ microspheres and indomethacin (1, 7.5 μg/larva) or PBS(control) and incubated in the dark at 30°C. Data represents the mean ± SE (*n* = 36, 432 in total across all four microsphere sizes). Symbol: * = P < 0.05 when comparing blank (negative controls) to indomethacin treatments at each respective time point. Unshared letters (a, b) represent significant differences (P ≤ 0.05) determined by Tukey’s multiple comparison tests.

Overall, significantly fewer 0.5 μm spheres made their way into the faeces within 24 hours and beyond (Figure 5d). This is likely due to the small size enabling movement across the gut barrier, and, those that remain within the midgut may get caught-up in tissue folds (Supp. Figure 2), thereby delaying passage through the hindgut/rectum. By analysing the control data (absent indomethacin) across the 72-hour period (Supp. Figure 3), we found a strong inverse correlation between microsphere size and haemolymph loads (Pearson Correlation; r = −0.988, P = 0.0115, R^2^ = 9.771) in support of our theory.

#### c) Gastric damage of the midgut

We further examined the effects of force-feeding indomethacin (7.5 μg/larva) and PBS (control) on *G. mellonella* larvae using wax (H&E) histology. The larval midgut in the presence of PBS did not show any clear symptoms of damage or altered tissue morphology (Figure 6 – upper panels). Uniform cellular arrangements of epithelial and goblet cells were observed, in addition to an intact basement membrane and visceral (striated) muscle layer. All control slides were considered grade 1 (0 – 2 discrete changes), with the exception of a single section at the 48-hour time point that was assigned grade 2 (Supp. Figure 4). Following indomethacin treatment, varying levels of tissue damage and midgut degradation were visible within 4 – 48 hours of all insects surveyed (Figure 6 – lower panels). Tissue slides were assigned damage grades 2 to 4 at 4 hours, and grades 2/3 at 24 and 48 hours (Supp. Figure 4) – representing at least five aberrations per slide (discrete, localised changes) to >50% compromised tissue (global damage; Figure 7). Deterioration of the larval gut manifested as sloughed epithelial cells, increased vacuolisation, partial/complete displacement of the gut lining into the lumen, cellular debris (or potential apoptotic bodies), membrane blistering/blebbing, nuclear condensation (pyknosis) and fragmentation (karyhorrhexis) (Figures 6 and 7). By 72 hours, the majority of slides were graded 1 and 2 (Supp. Figure 4), which suggests the tissue is being repaired.

**Figure 6a.**
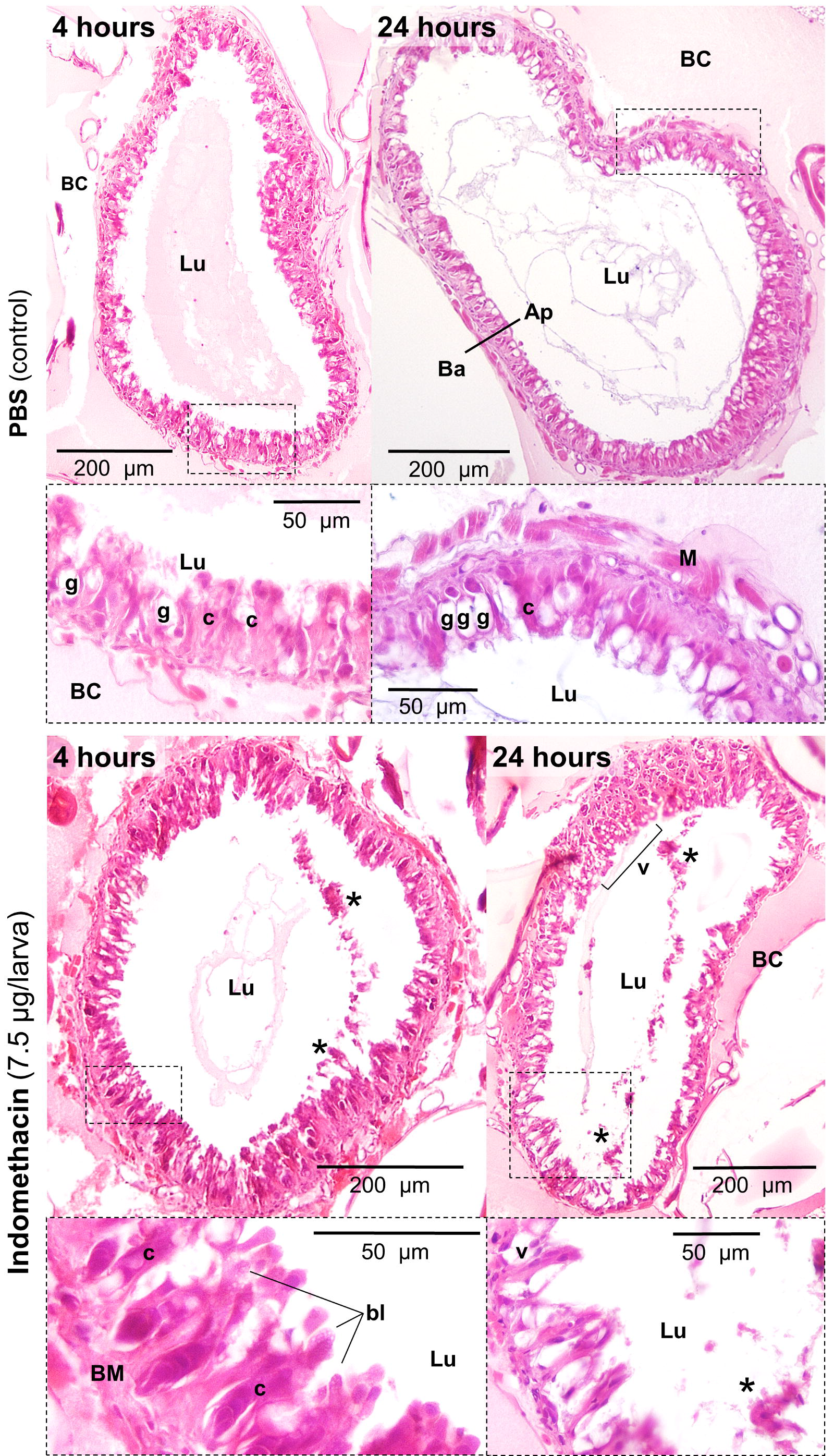
Gross histopathology of the midgut tissues from control and indomethacin-treated *Galleria mellonella*. Photomicrographs depict transverse sections of the midgut architecture at 4, 24, 48 and 72 hours after force-feeding PBS (control – upper panels) or indomethacin (7.5 μg – lower panels). Black boxes (broken lines) are used to highlight magnified regions of interest. Ap, apical; Ba, basolateral; BB, brush border; BC, body cavity; bl, blebbing/blistering of the cells; BM, basement membrane; c, columnar epithelial cell; g, goblet cell Lu, lumen; M, muscle; v, vacuole. An asterisk (*) denotes cellular damage and displacement into the lumen, and black arrows point to haemocytes (immune cells) within the body cavity (haemocoel).

**Figure 6b.**
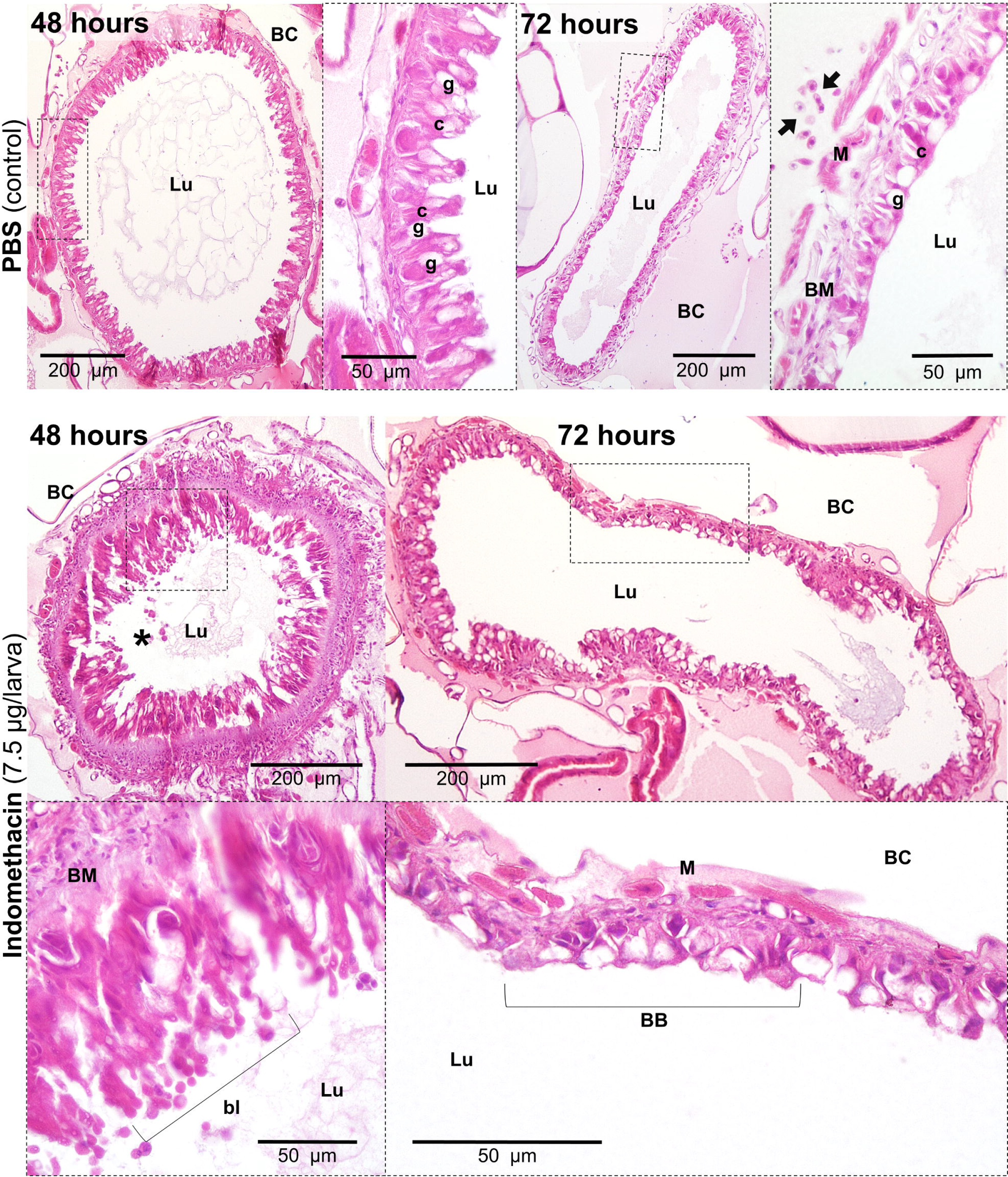

**Figure 7.**
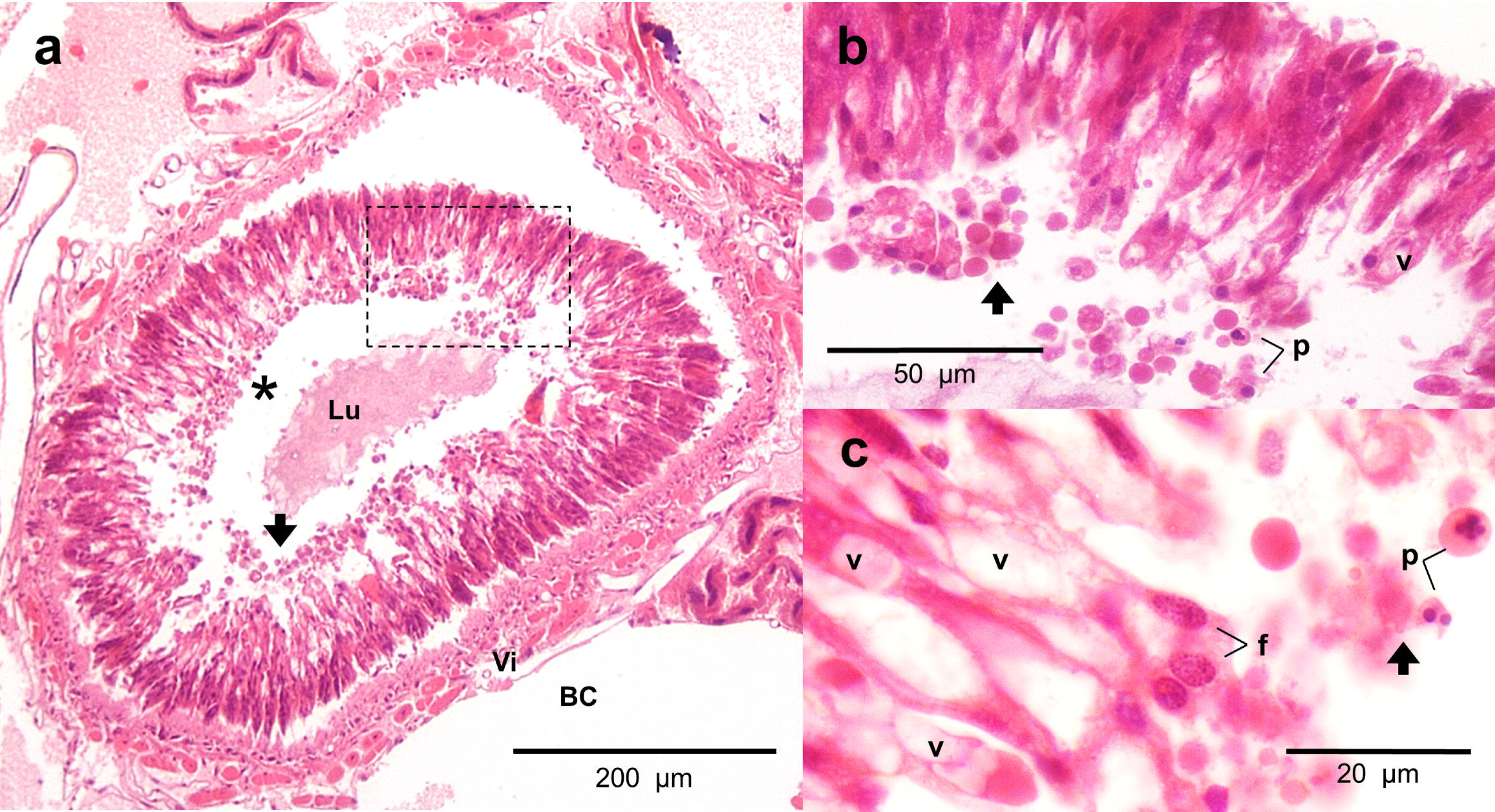
Histopathology of the midgut tissues from *Galleria mellonella* force-fed (7.5 μg) indomethacin. **a**) Transverse section of the larval midgut at 4 hours post-treatment. The photomicrograph displays severe global tissue damage, where almost the entire epithelium has dissociated (sloughed) from the basement membrane and visceral (Vi) muscle. The black arrows (all panels) signify the large number of cells displaced into the lumen (Lu). BC, body cavity (or haemocoel). **b & c)** These photomicrographs are magnified regions of the black box and asterisk, respectively. The epithelial cells are showing clear signs of death: nuclear pyknosis (p) and cytoplasmic fragmentation (f), distinct vacuolisation (v), and cell disintegration/debris (potential apoptotic bodies).

#### d) Gut repair (detoxification)

Administration of indomethacin by force feeding led to significant increases in glutathione-S-transferase (*F*(4, 40) = 7.35, P = 0.0002) and superoxide dismutase (F(4, 40) = 2.635 P = 0.0481) activites within the gut, but not in the haemolymph (GST; *F*(4, 40) = 0.5449, P = 0.704 and SOD; *F*(4, 40) = 0.801, P = 0.5.15; Figure 8). Larvae that were inoculated with 7.5 μg indoemthacin demonstrated consistently higher levels of GST activity in the gut, 2.3 – 3.2 Abs[340 nm]/min/mg, when compared to all other treatments/controls and timepoints (Figure 8a), whereas SOD activity peaked at 24 and 72 hours, ~0.4 Abs[560 nm]/min/mg (Figure 8c). A 2-way ANOVA revealed time to be a signifcant factor regarding SOD activity (*F*(3, 40) = 4.266, P = 0.0105), but this was not the case for GST (*F*(3, 40) = 2.584, P = 0.067). These patterns of enzyme acivity complement the restored nature of the midgut tissues at 72 hours in the histology (Figure 6). Detoxification-associated activities in the haemolypmh remained below 0.32 Abs[340 nm]/min/mg for GST and 0.15 Abs[560 nm]/min/mg for SOD (Figure 8b, 8d) – indicating strongly that the adverse effects of indomethacin were restricted to the gut.

**Figure 8.**
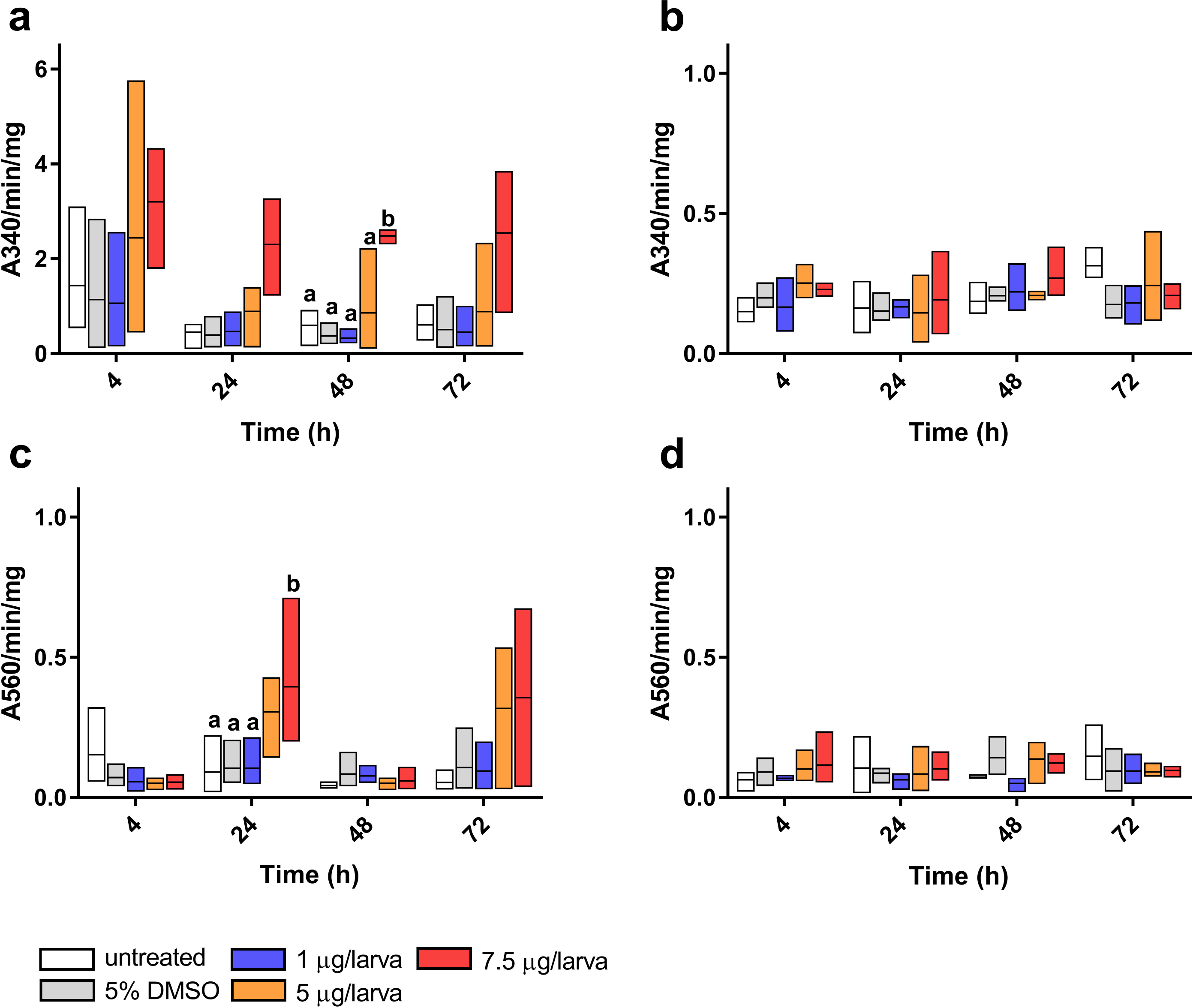
Detoxification-associated activities within *Galleria mellonella* following force-feeding of indomethacin, 0 – 7.5 μg/larva. Glutathione S-transferase activity was determined in the midgut (**a**) and haemolymph (**b**) by the change in 5-(2,4-dinitrophenyl)-glutathione accumulation (A340 nm). Superoxide dismutase activity was determined in the midgut (**c**) and haemolymph (**d**) via the inhibition of NBT reduction (A560 nm). Data are presented as floating bars (min, max) with mean lines shown (n = 36 per treatment, 180 in total; 3 insects were pooled at each time point). Unshared letters (a, b) represent statistical differences (P < 0.05) as determined by Tukey’s multiple comparison tests.

## Discussion

We assessed the relative toxicity of indomethacin in insect larvae via gavage (force-feeding) and intrahaemocoelic injection across a concentration range relevant to rodent models, 2–30 mg/kg (0.5–7.5 μg/larva), and found negligible side effects in terms of survival, development or immune cytotoxicity (Figures 1–3). When force-fed, larvae displayed broad symptoms of injury to the alimentary canal (Figures 5–8), e.g., a 3-fold increase in gut permeability and tissue degradation. We present considerable evidence that the integrity of the midgut is compromised by indomethacin within 4 to 24 hours, causing sufficient damage to activate repair/detoxification mechanisms (GST and SOD). The onset of indomethacin-associated gut leakiness has also been recorded within 24 hours in humans and mice, alongside ulceration and epithelial cell shrinkage (Playford *et al.*, 1999; 2001; Bjarnason and Takeuchi, 2009). Our combined use of X-ray microtomography and wax histology provide novel insight into the gross anatomy of the *G. mellonella* digestive tract – highlighting the suitability of the midgut for comparative pathobiology (Figures 4 – 7, Supp. Figure 2).

Indomethacin is a broad inhibitor of cyclooxygenase isozymes (COX1 and COX2), which are responsible for the initiation of prostaglandin synthesis (PGG_2_), and ultimately, the maintenance of inflammatory programmes and gastrointestinal mucosa (reviewed by Brune and Patrignani, 2015). In a previous study by Büyükgüzel *et al.* (2007), eicosanoid presence in the haemolymph was deemed essential for mediating nodule formation – a cellular defence reaction – during viremia. When they injected 50 μg of indomethacin into the haemocoel, there was a significant reduction in the number of nodules formed. We did not notice a reduction in the phagocytic capacity of haemocytes in insects force-fed 7.5 μg indomethacin and latex microspheres (Supp. Figure 1), which is also in contrast to earlier findings by Mandato *et al.* (1997). The authors reported on a reduction in the phagocytic index of *G. mellonella* haemocytes when exposed to 10μM indomethacin *in vitro* (Mandato *et al.*, 1997). A likely explanation for this discrepancy is that the majority of force-fed indomethacin remains in the midgut of our insects, despite the microspheres leaking into the surrounding haemolymph (where they are targeted by haemocytes; Supp. Figure 1). The immune cells of insects, namely the haemocytes, share many mechanistic and structural similarities with the innate immune cells of vertebrates, including pathogen recognition receptor signalling and phagocytosis-associated respiratory burst (Renwick *et al.*, 2007; Browne *et al.*, 2013; Butt *et al.*, 2016). Mandato *et al.* (1997) and Büyükgüzel *et al.* (2007) describe the immune-interference of indomethacin, and in addition to our observations of indomethacin-induced gut impairment, we consider that eicosanoid-like signalling should be added to the list of functional similairites between the innate immune systems of insects and vertebrates.

### Galleria mellonella *as an alternative animal model*

The fruit fly *Drosophila melanogaster* is a superior genetic resource, and over the last decade has been manipulated to gain novel insight into stem cell fate, immunity, antibiosis and homeostasis in the gut (Buchon *et al.*, 2009; Chandler *et al.*, 2011; Broderick and Lemaitre, 2012; Miguel-Aliaga *et al.*, 2018). Only recently, the genome of *G. mellonella* was made available (Lange *et al.*, 2018a), yet it stands unannotated. Furthermore, despite wax-moth larvae being used widespread as a screening tool for novel therapeutics, pathogenicity, and toxicology (reviewed by Tsai *et al.*, 2016), there remains a historical lag in molecular resources with the exception of some transcriptomic and miRNA data (Vogel *et al.*, 2011; Mukherjee and Vilcinskas, 2014). The financial and ethical incentives for using insect larvae over rodents and zebrafish are attractive, and so, we propose that vertebrates could be replaced partially for gut pathobiology. Waxmoth larvae are larger than traditional models like nematodes and drosophilids – this has two distinct advatages: (1) accurate doses can be administered orally (force-feeding), and (2) gram quanties of gut tissue can be obtained easily for downstream processing. Unlike other insect orders, the midgut of lepidopteran larvae represent the majority of tissue along the alimentary canal (mouth to anus; Figure 4), and, contains a specialised cell type, namely the goblet cell (Figure 6), which is also found in the human intestine (Engel and Moran, 2013; Linser and Dinglasan, 2014).

The most common larval inoculation technique is intrahaemocoelic injection of test compounds/microbes, however, force-feeding *G. mellonella* (i.e., gavage) is an emerging practice. When screening common food preservatives (e.g., potassium nitrate), Maguire *et al.* (2016) obtained comparable toxicology data (LD_50_) between insect larvae, human cell lines (HEp-2) and rats. We recently provided evidence that larvae can also be used to assess the lethality and putative immune-toxicological effects of shellfish poisoning toxins (e.g., okadaic acid) at FDA regulated levels in contaminated foods (Coates *et al.*, 2018). Okadaic acid that was force-fed to insects disrupted midgut homeostasis, leading to detrimental levels of lipid peroxidation (malondialdehyde accumulation) in a dose-dependent manner – resembling symptoms found often in the standardised mouse bioassay. Lange *et al.* (2018b) proved that *G. mellonella* could differentiate between an enteric symbiont (*Bacteroides vulgatus*) and pathobiont (*Escherichia coli*), and mount a strong immune response involving reaction oxygen/nitrogen species. Oral administration of the pathogen stimulated the up-regulation of immune-recognition genes in insects (e.g., apolipophorin III) and mice (e.g., Cd14). Interestingly, oral exposure of *G. mellonella* larvae to caffeine led to elevated levels of theobromine and theophylline in the haemolymph – suggesting that caffeine metabolism in this insect is similar to the process in mammals (Maguire *et al.*, 2017).

A better understanding of *G. mellonella*’s alimentary canal should assist insecticide development (e.g., boric acid and biopesticides, Büyükgüzel *et al.*, 2013; Grizanova *et al.*, 2014) as lepidopteran insects represent a sizeable number of devastating agricultural pests (e.g., *Spodoptera littoralis;* Linser and Dinglasan 2014). A key difference between the digestive systems of vertebrates and insects is the presence of phenoloxidase (PO) enzymes (Whitten and Coates, 2017). Phenoloxidases are responsible for the early processing of pigment precursors (quinones) into melanin, which plays several roles in development and immunity. Insect faeces are melanised upon release, and although the function of this is unclear, presumbaly it is due to gut phenoloxidase-activities and their oxidating/nitrosative by-products maintaining resident microbial populations from over-growing (Whitten and Coates, 2017). The gut microbiomes of insects are diverse and tend to be species specific – influenced invariably by diet and environmental factors (Engel and Moran, 2013). Few stuides have focussed on the *G. mellonella* gut microbiome, yet representatives of the Bacteroidetes, Firmicutes and Proteobacteria are homologous to the biota on microvilli of the human intestinal crypts (Mukherjee *et al.* 2013b; Dubovskiy *et al.*, 2016). Further work is needed to profile the residents of the insect gut, including fungi, viruses and Archaea. This is a timely topic, as wax-moth larvae are capable of degrading plastic (polypropylene; Bombelli *et al.*, 2017) – likely facilitated by the microbial consortium of the alimentary canal.

## Concluding remarks

We interrogated the physiological effects of indomethacin on *G. mellonella*, providing compelling evidence that indomethacin exposure leads to tissue damage, cell death, gut leakiness, and REDOX imbalance in insect larvae. This mimics closely the pathologIcal symptoms of their rodent counterparts. We describe the functional/structural similarities of lepidopteran midgut tissues to those found in regions of the human gastrointestinal tract. Our data reinforces the use of *G. mellonella* as a surrogate toxicology model, with a focus on screening neutraceuticals and food additives.

## Acknowledgements

We would like to thank Mrs Sophie Malkin (Bluefish Technical Officer, Swansea University) for her assistance with histology preparations, Dr Christopher B Cunningham (Swansea University) for helpful discussions, and Elizabeth Evans and Ria Mitchell (AIM Facility) for helpful tips with image data/segmentation.

## Author contributions

CJC conceived/designed the experiments. HE performed the experiments with assistance from CJC, AFR and RJ. HE collated the data. HE and CJC analysed and interpreted the data. CJC and HE prepared the manuscript, with input from AFR and RJ.

## References

Abràmoff, M. D., Magalhães, P. J., & Ram, S. J. (2004). Image processing with ImageJ. Biophotonics international, 11(7), 36–42.

Allegra, E., Titball, R. W., Carter, J., & Champion, O. L. (2018). *Galleria mellonella* larvae allow the discrimination of toxic and non-toxic chemicals. Chemosphere, 198, 469–472.

Altincicek, B., Linder, M., Linder, D., Preissner, K. T., & Vilcinskas, A. (2007). Microbial metalloproteinases mediate sensing of invading pathogens and activate innate immune responses in the lepidopteran model host *Galleria mellonella*. Infection and immunity, 75(1), 175–183.

Barnoy, S., Gancz, H., Zhu, Y., Honnold, C. L., Zurawski, D. V., & Venkatesan, M. M. (2017). The *Galleria mellonella* larvae as an in vivo model for evaluation of Shigella virulence. Gut microbes, 8(4), 335–350.

Basivireddy, J., Jacob, M., Ramamoorthy, P., Pulimood, A. B., & Balasubramanian, K. A. (2003). Indomethacin-induced free radical-mediated changes in the intestinal brush border membranes. Biochemical pharmacology, 65(4), 683–695.

Bjarnason, I., & Takeuchi, K. (2009). Intestinal permeability in the pathogenesis of NSAID-induced enteropathy. Journal of gastroenterology, 44(19), 23–29.

Bombelli, P., Howe, C. J., & Bertocchini, F. (2017). Polyethylene bio-degradation by caterpillars of the wax moth *Galleria mellonella*. Current Biology, 27(8), R292–R293.

Broderick, N. A., & Lemaitre, B. (2012). Gut-associated microbes of Drosophila melanogaster. Gut microbes, 3(4), 307–321.

Browne, N., Heelan, M., & Kavanagh, K. (2013). An analysis of the structural and functional similarities of insect hemocytes and mammalian phagocytes. Virulence, 4(7), 597–603.

Brune, K., & Patrignani, P. (2015). New insights into the use of currently available non-steroidal anti-inflammatory drugs. Journal of pain research, 8, 105.

Buchon, N., Broderick, N. A., Poidevin, M., Pradervand, S., & Lemaitre, B. (2009). Drosophila intestinal response to bacterial infection: activation of host defense and stem cell proliferation. Cell host & microbe, 5(2), 200–211.

Butt, T. M., Coates, C. J., Dubovskiy, I. M., & Ratcliffe, N. A. (2016). Entomopathogenic fungi: new insights into host–pathogen interactions. In Advances in genetics (Vol. 94, pp. 307–364). Academic Press.

Büyükgüzel, E., Büyükgüzel, K., Snela, M., Erdem, M., Radtke, K., Ziemnicki, K., & Adamski, Z. (2013). Effect of boric acid on antioxidant enzyme activity, lipid peroxidation, and ultrastructure of midgut and fat body of *Galleria mellonella*. Cell biology and toxicology, 29(2), 117–129.

Büyükgüzel, E., Tunaz, H., Stanley, D., & Büyükgüzel, K. (2007). Eicosanoids mediate *Galleria mellonella* cellular immune response to viral infection. Journal of insect physiology, 53(1), 99–105.

Campbell, P. M., Cao, A. T., Hines, E. R., East, P. D., & Gordon, K. H. (2008). Proteomic analysis of the peritrophic matrix from the gut of the caterpillar, *Helicoverpa armigera*. Insect biochemistry and molecular biology, 38(10), 950–958.

Champion, O. L., Wagley, S., & Titball, R. W. (2016). *Galleria mellonella* as a model host for microbiological and toxin research. Virulence, 7(7), 840–845.

Chandler, J. A., Lang, J. M., Bhatnagar, S., Eisen, J. A., & Kopp, A. (2011). Bacterial communities of diverse Drosophila species: ecological context of a host–microbe model system. PLoS genetics, 7(9), e1002272.

Card, R., Vaughan, K., Bagnall, M., Spiropoulos, J., Cooley, W., Strickland, T., … Anjum, M. F. (2016). Virulence characterisation of Salmonella enterica isolates of differing antimicrobial resistance recovered from UK livestock and imported meat samples. Frontiers in microbiology, 7, 640.

Coates, C. J., Lim, J., Harman, K., Rowley, A. F., Griffiths, D. J., Emery, H., & Layton, W. (2018). The insect, *Galleria mellonella*, is a compatible model for evaluating the toxicology of okadaic acid. Cell biology and toxicology, 1–14. DOI:10.1007/s10565-018-09448-2

Cools, F., Torfs, E., Aizawa, J., Vanhoutte, B., Maes, L., Caljon, G., … Cos, P. (2019). Optimization and Characterization of a *Galleria mellonella* Larval Infection Model for Virulence Studies and the Evaluation of Therapeutics Against *Streptococcus pneumoniae*. Frontiers in microbiology, 10.

Dubovskiy, I. M., Grizanova, E. V., Whitten, M. M., Mukherjee, K., Greig, C., Alikina, T., … Butt, T. M. (2016). Immuno-physiological adaptations confer wax moth Galleria mellonella resistance to Bacillus thuringiensis. Virulence, 7(8), 860–870.

Dubovskiy, I. M., Martemyanov, V. V., Vorontsova, Y. L., Rantala, M. J., Gryzanova,E. V., & Glupov, V. V. (2008). Effect of bacterial infection on antioxidant activity and lipid peroxidation in the midgut of Galleria mellonella L. larvae (Lepidoptera, Pyralidae). Comparative Biochemistry and Physiology Part C: Toxicology & Pharmacology, 148(1), 1–5.

Engel, P., & Moran, N. A. (2013). The gut microbiota of insects–diversity in structure and function. FEMS microbiology reviews, 37(5), 699–735.

Green, L. F., Bergquist, P. R., & Bullivant, S. (1980). The structure and function of the smooth septate junction in a transporting epithelium: the Malpighian tubules of the New Zealand glow-worm *Arachnocampa luminosa*. Tissue and Cell, 12(2), 365–381.

Grizanova, E. V., Dubovskiy, I. M., Whitten, M. M. A., & Glupov, V. V. (2014). Contributions of cellular and humoral immunity of Galleria mellonella larvae in defence against oral infection by *Bacillus thuringiensis*. Journal of invertebrate pathology, 119, 40–46.

Kloezen, W., van Helvert-van Poppel, M., Fahal, A. H., & van de Sande, W. W. (2015). A Madurella mycetomatis grain model in Galleria mellonella larvae. PLoS neglected tropical diseases, 9(7), e0003926.

Kuraishi, T., Binggeli, O., Opota, O., Buchon, N., & Lemaitre, B. (2011). Genetic evidence for a protective role of the peritrophic matrix against intestinal bacterial infection in *Drosophila melanogaster*. Proceedings of the National Academy of Sciences, 108(38), 15966–15971.

Lange, A., Beier, S., Huson, D. H., Parusel, R., Iglauer, F., & Frick, J. S. (2018a). Genome sequence of *Galleria mellonella* (greater wax moth). Genome Announc., 6(2), e01220–17.

Lange, A., Schäfer, A., Bender, A., Steimle, A., Beier, S., Parusel, R., & Frick, J. S. (2018b). Galleria mellonella: a novel invertebrate model to distinguish intestinal symbionts from pathobionts. Frontiers in immunology, 9.

Lim, J., Coates, C. J., Seoane, P. I., Garelnabi, M., Taylor-Smith, L. M., Monteith, P., … May, R. C. (2018). Characterizing the Mechanisms of Nonopsonic Uptake of Cryptococci by Macrophages. The Journal of Immunology, ji1700790.

Linser, P. J., & Dinglasan, R. R. (2014). Insect gut structure, function, development and target of biological toxins. In Advances in insect physiology (Vol. 47, pp. 1–37). Academic Press.

Limaye, A. (2012). Drishti: a volume exploration and presentation tool. In Developments in X-Ray Tomography VIII. International Society for Optics and Photonics (Vol. 8506, p. 85060X).

Miguel-Aliaga, I., Jasper, H., & Lemaitre, B. (2018). Anatomy and Physiology of the Digestive Tract of Drosophila melanogaster. Genetics, 210(2), 357–396.

Maguire, R., Duggan, O., & Kavanagh, K. (2016). Evaluation of Galleria mellonella larvae as an in vivo model for assessing the relative toxicity of food preservative agents. Cell biology and toxicology, 32(3), 209–216.

Maguire, R., Kunc, M., Hyrsl, P., & Kavanagh, K. (2017). Caffeine administration alters the behaviour and development of Galleria mellonella larvae. Neurotoxicology and teratology, 64, 37–44.

Mahmood, A., Fitzgerald, A. J., Marchbank, T., Ntatsaki, E., Murray, D., Ghosh, S., & Playford, R. J. (2007). Zinc carnosine, a health food supplement that stabilises small bowel integrity and stimulates gut repair processes. Gut, 56(2), 168–175.

Mandato, C. A., Diehl-Jones, W. L., Moore, S. J., & Downer, R. G. (1997). The effects of eicosanoid biosynthesis inhibitors on prophenoloxidase activation, phagocytosis and cell spreading in Galleria mellonella. Journal of Insect Physiology, 43(1), 1–8.

Marchbank, T., Ojobo, E., Playford, C. J., & Playford, R. J. (2011). Reparative properties of the traditional Chinese medicine *Cordyceps sinensis* (Chinese caterpillar mushroom) using HT29 cell culture and rat gastric damage models of injury. British journal of nutrition, 105(9), 1303–1310.

Matsui, H., Shimokawa, O., Kaneko, T., Nagano, Y., Rai, K., & Hyodo, I. (2011). The pathophysiology of non-steroidal anti-inflammatory drug (NSAID)-induced mucosal injuries in stomach and small intestine. Journal of clinical biochemistry and nutrition, 48(2), 107–111.

Mowlds, P., Coates, C., Renwick, J., & Kavanagh, K. (2010). Dose-dependent cellular and humoral responses in *Galleria mellonella* larvae following β-glucan inoculation. Microbes and infection, 12(2), 146–153.

Mukherjee, K., Hain, T., Fischer, R., Chakraborty, T., & Vilcinskas, A. (2013a). Brain infection and activation of neuronal repair mechanisms by the human pathogen *Listeria monocytogenes* in the lepidopteran model host *Galleria mellonella*. Virulence, 4(4), 324–332.

Mukherjee, K., Raju, R., Fischer, R., & Vilcinskas, A. (2013b). *Galleria mellonella* as a model host to study gut microbe homeostasis and brain infection by the human pathogen Listeria monocytogenes. In Yellow Biotechnology I (pp. 27–39). Springer, Berlin, Heidelberg

Mukherjee, K., & Vilcinskas, A. (2014). Development and immunity-related microRNAs of the lepidopteran model host Galleria mellonella. BMC genomics, 15(1), 705.

Perron, N., Tremblay, E., Ferretti, E., Babakissa, C., Seidman, E. G., Levy, E., … Beaulieu, J. F. (2013). Deleterious effects of indomethacin in the mid-gestation human intestine. Genomics, 101(3), 171–177.

Playford, R. J., Floyd, D. N., Macdonald, C. E., Calnan, D. P., Adenekan, R. O., Johnson, W., … Marchbank, T. (1999). Bovine colostrum is a health food supplement which prevents NSAID induced gut damage. Gut, 44(5), 653–658.

Playford, R. J., Macdonald, C. E., Calnan, D. P., Floyd, D. N., Podas, T., Johnson, W., … Marchbank, T. (2001). Co-administration of the health food supplement, bovine colostrum, reduces the acute non-steroidal anti-inflammatory drug-induced increase in intestinal permeability. Clinical Science, 100(6), 627–633.

Renwick, J., Reeves, E. P., Wientjes, F. B., & Kavanagh, K. (2007). Translocation of proteins homologous to human neutrophil p47phox and p67phox to the cell membrane in activated hemocytes of Galleria mellonella. Developmental & Comparative Immunology, 31(4), 347–359.

Senior, N. J., Bagnall, M. C., Champion, O. L., Reynolds, S. E., La Ragione, R. M., Woodward, M. J., … Titball, R. W. (2011). Galleria mellonella as an infection model for *Campylobacter jejuni* virulence. Journal of medical microbiology, 60(5), 661–669.

Smecuol, E., Bai, J. C., Sugai, E., Vazquez, H., Niveloni, S., Pedreira, S., … Meddings, J. (2001). Acute gastrointestinal permeability responses to different non-steroidal anti-inflammatory drugs. Gut, 49(5), 650–655.

Sigthorsson, G., Crane, R., Simon, T., Hoover, M., Quan, H., Bolognese, J., & Bjarnason, I. (2000). COX-2 inhibition with rofecoxib does not increase intestinal permeability in healthy subjects: a double blind crossover study comparing rofecoxib with placebo and indomethacin. Gut, 47(4), 527–532.

Strober, W. (2015). Trypan blue exclusion test of cell viability. Current protocols in immunology, 111(1), A3–B.

Tsai, C. J. Y., Loh, J. M. S., & Proft, T. (2016). *Galleria mellonella* infection models for the study of bacterial diseases and for antimicrobial drug testing. Virulence, 7(3), 214–229.

Vogel, H., Altincicek, B., Glöckner, G., & Vilcinskas, A. (2011). A comprehensive transcriptome and immune-gene repertoire of the lepidopteran model host Galleria mellonella. BMC genomics, 12(1), 308.

Wagley, S., Borne, R., Harrison, J., Baker-Austin, C., Ottaviani, D., Leoni, F., … Titball, R. W. (2018). Galleria mellonella as an infection model to investigate virulence of Vibrio parahaemolyticus. Virulence, 9(1), 197–207.

Whitten, M. M., & Coates, C. J. (2017). Re‐evaluation of insect melanogenesis research: Views from the dark side. Pigment cell & melanoma research, 30(4), 386–401.

