## Supplemental Information for "Indomethacin-induced gut damage in a surrogate insect model, *Galleria mellonella*"

*Corresponding author:

**Supplementary Materials/Methods:**

*Cole’s hematoxylin and eosin staining*

Slides were dewaxed using Histochoice/Histoclear for 15 mins prior to washing in absolute ethanol for 2 mins, followed by 1 min in 90% ethanol and 1 min in 70% ethanol before staining in Cole’s haematoxylin for 13 mins. Once stained in haematoxylin, slides were then washed in Scotts’ solutions for 2 mins to stain nuclear material blue. Samples were washed in water for 10 seconds to remove any excess stain. At this point, several slides were selected and viewed under a microscope to check the staining before proceeding with Eosin. Samples were dehydrated in 70% ethanol for 1 min, followed by Eosin for 6 mins. Once stained in Eosin, samples were dipped in 70% ethanol for a few seconds to remove excess stain. Slides were viewed under a microscope at this point to check the stain was sufficient. Slides were further dehydrated in 90% ethanol for 10 seconds and then in absolute ethanol for 5 mins twice. Finally, slides were washed three times in fresh Histochoice/Histoclear each for 5 mins three times prior to coverslip mounting using DPX. Mounted slides were trasnfered to a heating rack for 10 mins before being placed in an incubator at 40 °C until dry, and the DPX had set.

*Inspection of histology slides*

A grading system (1 – 4) was used to indicate visible damage or lack thereof by modifying the guidelines developed by *Watanabe et al*. (2017):

1) discrete change with no more than 0 – 2 tissue aberrations per slide

2) discrete changes of 3 to 5 tissue aberrations per slide

3) spatial change representing ≥25% damage (the alteration is dramatic)

4) global change of >50% of a specific tissue type or the entire slide.

Watanabe, H., Horie, Y., Takanobu, H., Koshio, M., Flynn, K., Iguchi, T., & Tatarazako, N. (2017). Medaka extended one‐generation reproduction test evaluating 4‐nonylphenol. *Environmental toxicology and chemistry*, *36*(12), 3254-3266.

<http://www.oecd.org/env/ehs/testing/Histopath-Guidance-Document-for-the-Medeka-Extended-One-Generation-Reproduction-Test-Part_1.pdf>

**Supplementary results**:

**Supplementary Table 1** Proportions of haemocytes staining positively for Trypan-blue (i.e., cell death) when treated with indomethacin (1 – 7.5 μg/larva) or PBS.

|  | 4 hours | 24 hours | 48 hours | 72 hours |
| --- | --- | --- | --- | --- |
| untreated | 9.9% | 8.3% | 9% | 10.2% |
| I**njection** |  |  |  |  |
| PBS + 5% DMSO | 8.5% | 8% | 10.6% | 10.3% |
| 1 μg/larva | 9.8% | 10.7% | 12.1% | 10.5% |
| 5 μg/larva | 9.2% | 10.7% | 9.4% | 11% |
| 7.5 μg/larva | 10.8% | 11.9% | 11.9% | 8.8% |
| **Force feeding** |  |  |  |  |
| PBS + 5% DMSO | 11.3% | 9% | 8.1% | 9.4% |
| 1 μg/larva | 10.8% | 7.2% | 10.8% | 10.3% |
| 5 μg/larva | 8.9% | 8.5% | 9.2% | 9% |
| 7.5 μg/larva | 7.1% | 8.6% | 9.3% | 9.2% |

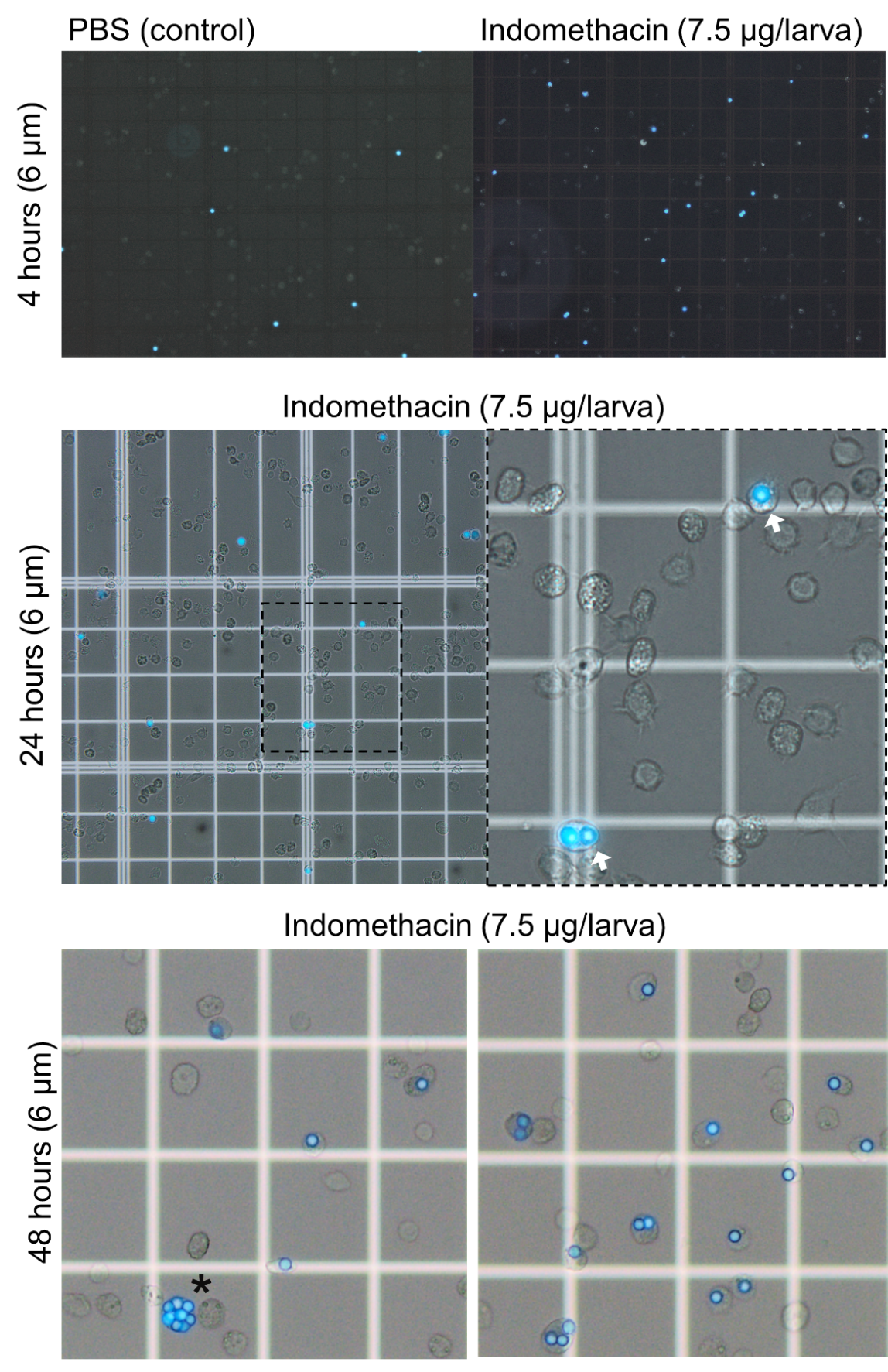

**Supplementary Figure 1. Latex microspheres (6 µm) in the haemolymph of *Galleria mellonella* co-inoculated with indomethacin (7.5 µg/larva).** The top, middle and bottom panels represent images taken from 4, 24 and 48-hours post-inoculation, respectively. The white arrows point to spheres within the cytoplasm of phagocytic haemocytes (immune cells).The asterisk (*) signifies a haemocyte with eight internalised microspheres.

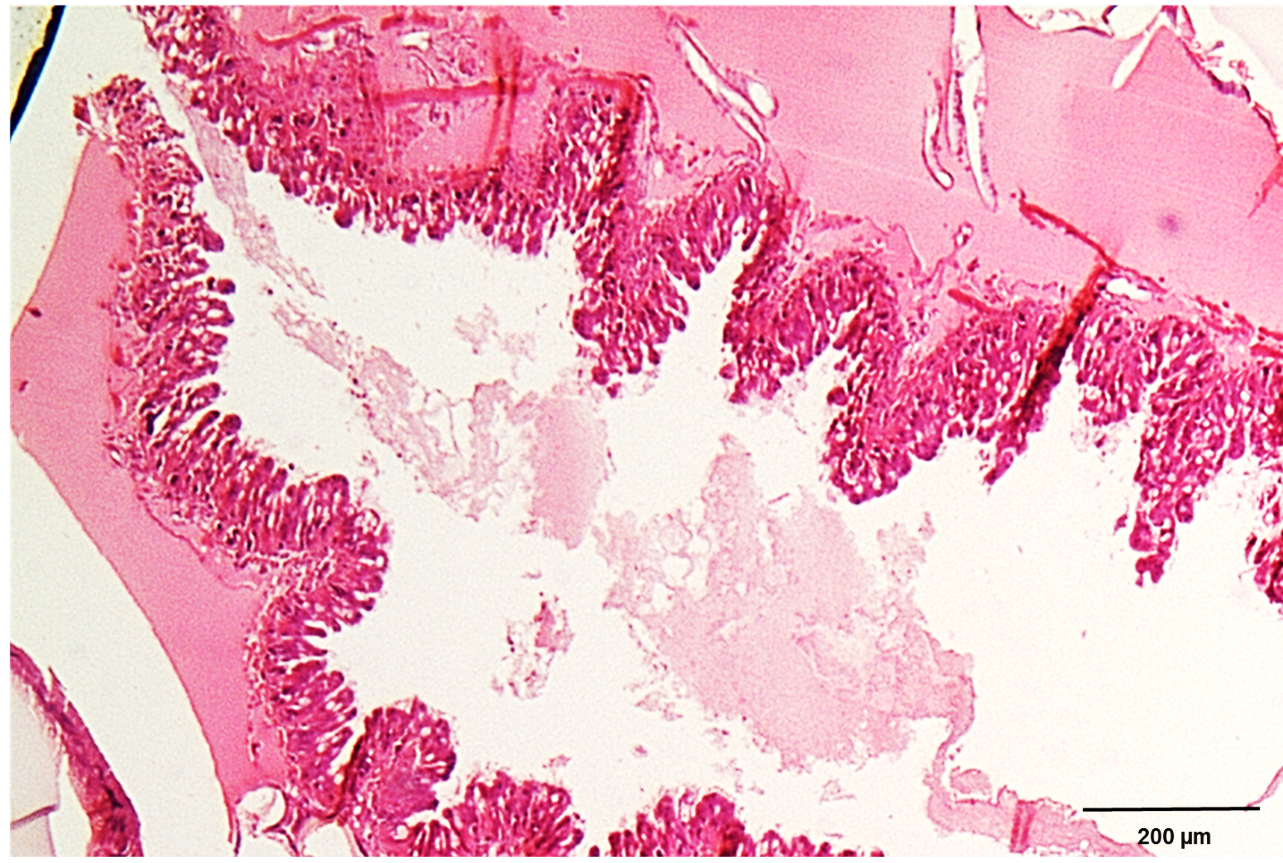

**Supplementary Figure 2.** **Histology (longitudinal) section of the *Galleria mellonella* midgut**. The photomicrograph depicts the series of folds/pleats that make-up the midgut.

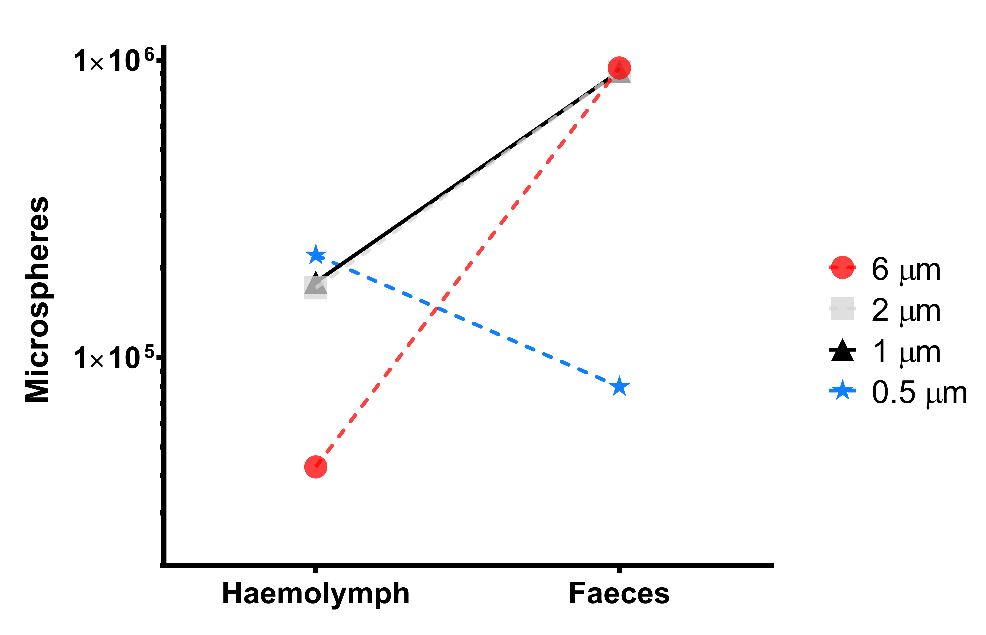

**Supplementary Figure 3. Gut permeability of *Galleria mellonella* larvae following force feeding of microspheres (0.5 – 6 µm).** The data presented are from the control (no indomethacin) larvae only over the entire experimental period (72 hours), i.e., the total microspheres recovered. There is an inverse correletion between microsphere diameter and haemolypmh load – the larger a microsphere is the less likely it will make its way through the gut wall and into the body cavity (haemocoel).

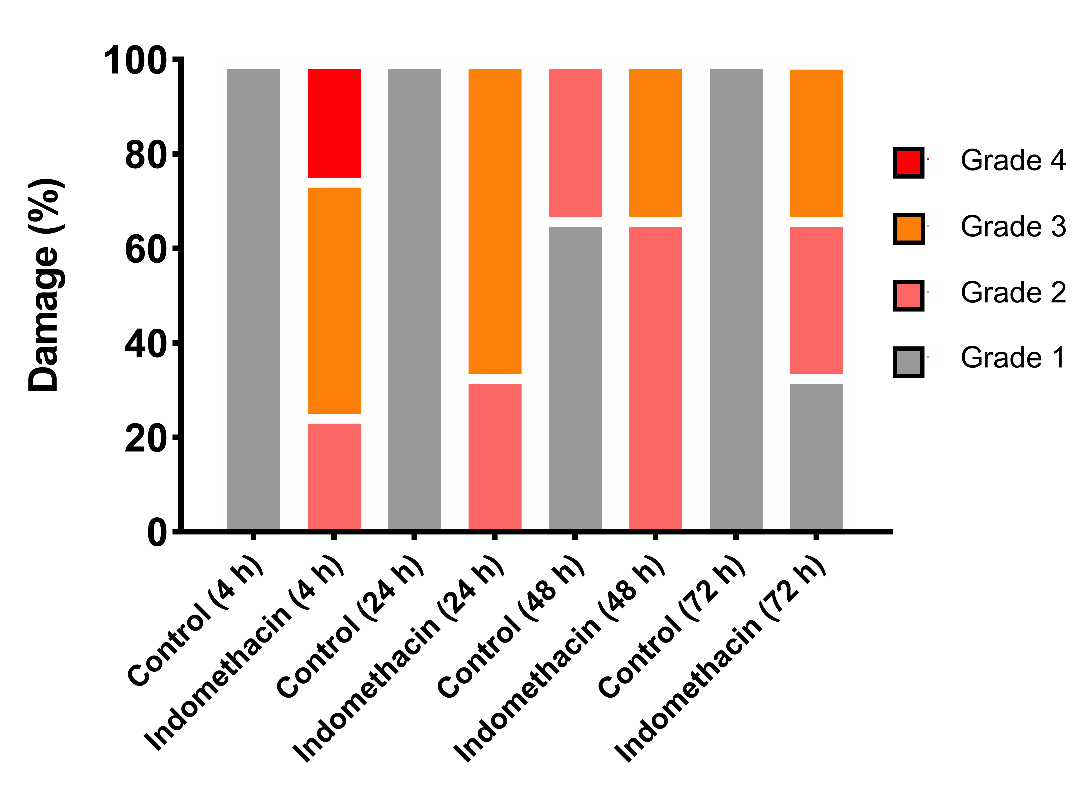

**Supplementary Figure 4. The extent of midgut tissue damage in *Galleria mellonella* force-fed indomethacin (7.5 µg/larva) or PBS.** Histology slides were single blind assessed in pairs (treatment vs control) and subsequently assigned a grade (1 - 4) based on damage(s). Grade 1 indicates little to no damage, whereas Grade 4 represents global damage affecting >50% of tissue. Data have been compiled from assessments carried out at 4 (n = 8), 24 (n = 6), 48 (n = 6) and 72 (n = 6) hours post-inoculation.
